# A hypermorphic *Nfkbid* allele represents an *Idd7* locus gene contributing to impaired thymic deletion of autoreactive diabetogenic CD8^+^ T-cells in NOD mice

**DOI:** 10.1101/249094

**Authors:** Maximiliano Presa, Jeremy J. Racine, Jennifer R. Dwyer, Deanna J. Lamont, Jeremy J. Ratiu, Vishal Kumar Sarsani, Yi-Guang Chen, Aron Geurts, Ingo Schmitz, Timothy Stearns, Jennifer Allocco, Harold D. Chapman, David V. Serreze

## Abstract

In both NOD mice and humans, the development of type 1 diabetes (T1D) is dependent in part on autoreactive CD8^+^ T-cells recognizing pancreatic ß-cell peptides presented by often quite common MHC class I variants. Studies in NOD mice previously revealed the common H2-K^d^ and/or H2-D^b^ class I molecules expressed by this strain acquire an aberrant ability to mediate pathogenic CD8^+^ T-cell responses through interactions with T1D susceptibility (*Idd*) genes outside the MHC. A gene(s) mapping to the *Idd7* locus on proximal Chromosome 7 was previously shown to be an important contributor to the failure of the common class I molecules expressed by NOD mice to mediate the normal thymic negative selection of diabetogenic CD8^+^ T-cells. Using an inducible model of thymic negative selection and mRNA transcript analyses we initially identified an elevated *Nfkbid* expression variant is likely an NOD *Idd7* region gene contributing to impaired thymic deletion of diabetogenic CD8^+^ T-cells. CRISPR/Cas9-mediated genetic attenuation of *Nfkbid* expression in NOD mice resulted in improved negative selection of autoreactive diabetogenic AI4 and NY8.3 CD8^+^ T-cells. These results indicated allelic variants of *Nfkbid* represent an *Idd7* gene contributing to the efficiency of intrathymic deletion of diabetogenic CD8^+^ T-cells. However, while enhancing thymic deletion of pathogenic CD8^+^ T-cells, ablation of *Nfkbid* expression surprisingly accelerated T1D onset in NOD mice likely at least in part by numerically decreasing regulatory T- and B-lymphocytes (Tregs/Bregs), thereby reducing their peripheral immunosuppressive effects.

## Introduction

In both the NOD mouse model and humans, type 1 diabetes (T1D^3^) results from the autoimmune destruction of insulin-producing pancreatic β-cells mediated by the combined activity of CD4^+^ and CD8^+^ T-cells, as well as B-lymphocytes (1). T1D is highly polygenic in nature, as more than 50 loci have been associated with disease susceptibility or resistance in both humans and NOD mice (2, 3). However, while its complete pathogenic etiology remains unsolved, there is a wide acceptance that disease develops as a consequence of interactions between T1D susceptibility (*Idd*) genes resulting in breakdowns in mechanisms controlling induction of immunological tolerance to self-proteins (4, 5). Some *Idd* genes appear to contribute to defects in central tolerance mechanisms that normally represent a first checkpoint engendering the thymic deletion of autoreactive CD4^+^ and CD8^+^ T-cells during early stages of their development (6–8). However, even in healthy individuals, some autoreactive T-cells escape central tolerance mechanisms (9). Such autoreactive effectors are normally prevented from mediating pathogenic effects by a second checkpoint, provided by multiple mechanisms of peripheral tolerance, with significant evidence for some of these processes also being disrupted by various *Idd* genes (5).

In the thymus, immature CD4^+^CD8^+^ double positive (DP) thymocytes are able to recognize self-peptide-MHC complexes displayed on antigen presenting cells (APC, thymic epithelial cells, dendritic cells (DC) or macrophages) through interactions with clonally distributed T-cell receptor (TCR) molecules (10). The strength of TCR signaling resulting from this TCR-peptide-MHC interaction dictates the fate of immature DP thymocytes (9, 10). DP thymocytes undergoing high avidity interactions with self-peptide-MHC complexes are normally deleted by apoptosis or diverted to a CD4^+^Foxp3^+^ regulatory T-cell lineage (Treg) (9). In both humans and NOD mice, defects in central tolerance occur at three levels: reduced presentation of self-antigen in thymus; defects in post-TCR engagement signaling events; and variability in the basal or induced expression level of molecules controlling the final pro-/anti-apoptotic balance (4, 5). Deficiencies in this process contribute to the thymic survival and peripheral seeding of autoreactive T-cells that overwhelm the suppressive capacity of self-antigen-specific Tregs (5). Several *Idd* genes and loci have been identified as contributing to such central tolerance induction defects (4, 5). Among these are MHC haplotypes long known to provide the primary T1D genetic risk factor (11). Within the MHC, certain unusual class II molecules play an important role in T1D pathogenesis by allowing the development and functional activation of autoreactive CD4^+^ T-cells (12). However, MHC class II expression is largely restricted to thymic epithelial cells and hematopoietically derived APC (10, 13). Thus, while class II restricted autoreactive CD4^+^ T-cells clearly play a critical role in T1D development, they cannot do so by directly engaging insulin-producing pancreatic β-cells (14). Conversely, like most other tissues, pancreatic β-cells do express MHC class I molecules. Hence, through an ability to directly engage and destroy pancreatic β cells, MHC class I restricted autoreactive CD8^+^ T-cells are likely the ultimate mediators of T1D development in both humans and NOD mice (14, 15). Epidemiological studies have shown that in addition to class II effects, particular HLA class I variants provide an independent T1D risk factor in humans (16, 17). Interestingly, this includes some quite common MHC class I variants that in both humans and NOD mice only acquire an aberrant ability to contribute to T1D when expressed in the context of other *Idd* genes.

Previous studies found interactive contributions from some polymorphic *Idd* genes outside of the MHC determine the extent to which particular class I molecules can allow the development and functional activation of diabetogenic CD8^+^ T-cells. This was initially demonstrated by congenically transferring a transgenic Vα8^+^Vβ2^+^ TCR from the H2^g7^ class I restricted diabetogenic CD8^+^ T-cell clone AI4 from a NOD genetic background strain (NOD-AI4) to a T1D resistant C57BL/6-stock (B6) congenic for the *H2^g7^* haplotype (B6.*H2^g7^*). Autoreactive AI4 TCR transgenic T-cells were deleted to a significantly greater extent at the DP stage of thymic development in B6.*H2^g7^* than NOD mice (7). This result indicated non-MHC genes allelically differing in NOD and B6 mice regulate the extent to which autoreactive diabetogenic CD8^+^ T-cells undergo thymic deletion. Such non-MHC genes controlling differing thymic negative selection efficiency in NOD and B6 background mice were found to function in a T-cell intrinsic manner (7). Subsequent linkage analyses and congenic strain evaluation identified a region on proximal Chromosome (Chr.) 7 to which the *Idd7* locus had been previously mapped (18, 19), as containing a gene strongly contributing to the differential thymic deletion efficiency of diabetogenic AI4 CD8^+^ T-cells in NOD and B6.*H2^g7^* mice (8).

In this study, we identify differential expression level variants of *Nfkbid* as a likely *Idd7* region gene regulating the efficiency of diabetogenic CD8^+^ T-cell thymic negative selection. *Nfkbid* (aka *I*κ*BNS*) is a family member of the non-conventional IκB modulators of the transcription factor NF-κB (20). Depending on cell type and physiological context, *Nfkbid* can either positively or negatively modulate NF-κB activity (20). *Nfkbid* was first identified as a gene expressed in thymocytes undergoing negative, but not positive selection upon antigen specific TCR stimulation (21). Curiously, genetic ablation of *Nfkbid* in B6 background mice did not elicit any alteration in thymocyte subset distribution or any overt signs of autoimmunity (22). Using TCR transgenic mouse models to analyze antigen specific thymic negative selection, we show in this report that a higher expression variant of *Nfkbid* in NOD (compared to B6 mice) contributes to diminished negative selection in the former strain of both the AI4 and NY8.3 MHC class I restricted autoreactive diabetogenic CD8^+^ T-cell clonotypes. While genetic attenuation of *Nfkbid* expression to levels similar to that in B6 background mice results in improved thymic deletion of pathogenic CD8^+^ T-cells, total ablation of this gene also diminishes levels of peripheral Tregs and regulatory B-lymphocytes (Bregs) resulting in accelerated T1D development in NOD mice.

## Materials and Methods

### Mouse strains

NOD/ShiLtDvs (hereafter NOD), C57BL/6J (B6), and NOD.Cg-*Prkdc^scid^* Emv30b/Dvs (NOD.*scid*) mice are maintained in a specific pathogen free research colony at The Jackson Laboratory. NOD.129X1(Cg)-Foxp3^tm2Tch^/Dvs (hereafter NOD.*Foxp3-eGFP*), carrying the reporter construct *Foxp3^tm2(eGFP)Tch^*, were previously described (23, 24). NOD mice carrying a transgenic Vα8.3^+^Vβ2^+^ TCR derived from the AI4 diabetogenic CD8^+^ T-cell clone (NOD-AI4) have also been previously described (25). Previously generated NOD.LCMV mice transgenically express a Vα2^+^Vβ8^+^ TCR from a CD8^+^ T-cell clone recognizing the H2-D^b^ restricted gp33 peptide (KAVYNFATM) derived from Lymphocytic Choriomeningitis Virus (LCMV) (26, 27). NOD.NY8.3 mice transgenically expressing the TCR derived from the H2-K^d^ restricted diabetogenic NY8.3 CD8^+^ T-cell clone (28) are maintained in an heterozygous state. A NOD stock congenic for a segment of B6-derived Chr. 7 delineated by the flanking markers *Gpi^b^* and *D7Mit346* (29) (officially designated NOD.B6-(*Gpi1-D7Mit346*)/LtJ and hereafter abbreviated NOD.*Chr7^B6^*FL) was used to introduce the congenic interval into NOD-AI4 mice (8).

### Congenic truncation analysis

The NOD.*Chr7^B6^*FL stock was intercrossed with NOD-AI4. Resultant F1 progeny were intercrossed and all F2 offspring were analyzed for recombination events within the original Chr. 7 congenic region delineated by the flanking markers *Gpi^b^* (34.2 Mb) and *D7Mit346* (58.7 Mb). Genomic tail DNA samples were screened for the microsatellite markers *D7Mit267*, *D7Mit117, D7Mit155, Gpi1^b^, D7Mit79, D7Mit225, D7Mit78, D7Mit247, D7Mit270, D7Mit230,* and *D7Mit346* by PCR. Identified recombinant mice were backcrossed to NOD-AI4 and then intercrossed to generate progeny homozygous for the sub-congenic regions. The homozygous state for each sub-congenic line was verified by PCR analyses of the markers listed above. Two lines resulted from this process, Ln82 with a sub-congenic region *D7Mit117*–*D7Mit247* and Ln16, with a sub-congenic region *D7Mit79*–*D7Mit247*.

### Generation of NOD Nfkbid-deficient mice

CRISPR/Cas9 technology was utilized to directly ablate the *Nfkbid* gene in NOD mice. NOD/ShiLtDvs embryos were microinjected with 3pl of a solution containing Cas9 mRNA and single guide RNA (sgRNA) at respective concentrations of 100ng/µl and 50ng/µl. The sgRNA sequence (5’-AGGCCCATTTCCCCTGGTGA-3’) was designed to delete the transcriptional start site of *Nfkbid* in exon 3. Genomic tail DNA was screened by targeted Sanger sequencing: the genomic region around exon 3 was amplified by PCR with primers Nfkbid-KO-F1 (5’-TGCTTGAGATCCAGTAG-3’) and Nfkbid-KO-R1 (5’-CCCTGACATCTCAGAATA-3’). The resulting PCR product was purified and sequenced using an ABI 3730 DNA analyzer (Applied Biosystems-Thermo Fisher Scientific, Waltham, MA, USA). Analyses of Sanger sequencing data was done using the software Poly Peak Parser (30) which allows identification of the different alleles present in heterozygous mutant mice. During the screening of N1 mutants, we identified an allele characterized by an 8-nucleotide deletion and an insertion of 154-nucleotides originated by copy and inversion of a DNA segment from *Nfkbid* exon 2. This produced a major disruption of the *Nfkbid* gene and deletion of the reference translation start site (Supplementary Fig. 1). N1F1 NOD-*Nfkbid^+/−^*mutant mice were intercrossed and the disrupted allele fixed to homozygosity. The new stock, formally designated NOD/ShiLtDvs-*Nfkbid*^<EM3DVS>/^Dvs (hereafter abbreviated NOD-*Nfkbid^−/−^*), was maintained by brother-sister matting.

Previous reports indicated that *Nfkbid*-deficient mice have a diminished Foxp3^+^ Treg compartment (31). Thus, we introduced the *Foxp3^tm2(eGFP)Tch^* reporter construct previously congenically transferred to the NOD strain (24) into the NOD-*Nfkbid^−/−^* stock which was subsequently fixed to homozygosity.

### Nfkbid gene expression analysis

Total RNA from whole thymus or sorted DP cell lysates was extracted using the RNeasy Mini kit (Qiagen, Germantown, MD, USA) following the vendor instructions. For whole thymus tissue RNA, the complete organ was put in 2ml of RNAlater stabilization solution (Thermo Fisher Scientific), incubated 24 h at room temperature and then stored at -20°C until processing. In other experiments 2×10^5^ DP cells were sorted in 100% FBS, washed 2 times in cold PBS and resuspended in RLT lysis buffer (Qiagen) and stored at -20°C until processing. Extracted total RNA was quantified by NanoDrop (Thermo Fisher Scientific) and quality assessed in Bioanalyzer (Agilent Technologies, Santa Clara, CA, USA). Samples with a RNA integrity index (RIN index) of 7 or higher were used for gene expression analyses. For cDNA synthesis 500ng of total RNA was diluted to 5μl in DEPC treated water and mixed with 5μl of SuperScript IV VILO Master Mix (Thermo Fisher Scientific) following vendor instructions.

Expression of *Nfkbid* was assessed by a predesigned and validated TaqMan qPCR assay using Mm.PT.58.12759232 (IDT Integrated Technologies, Coralville, Iowa, USA) targeting the exon 3-4 region and able to identify all four possible transcripts from wild type alleles. The assay consists of a FAM-labeled and double-quenched probe (5’-/56-FAM-CCTGGTGAT-/ZEN/-GGAGGACTCTCTGGAT-/3IABkFQ/-3’) that spans the 8-nucleotide deletion site and the primers p1 (5’-GACAGGGAAGGCTCAGGATA-3’) and p2 (5’-GCTTCCTGACTCCTGATTTCTAC-3’). The assay was used at a final concentration of 500nM primers and 250nM probe. As an endogenous control, we used a predesigned and validated TaqMan assay for *Gapdh* (ID: Mm99999915_g1) (VIC-labeled) (Applied Biosystems – Thermo Fisher Scientifc). The qPCR analyses were done following the manufacturer’s protocol using a ViiA-7 real time PCR system (Applied Biosystems - Thermo Fisher Scientific). Relative gene expression was determined by the ddCt method using the application RQ in Thermo Fisher Cloud Software, Version 1.0 (Thermo Fisher Scientific).

### Microarray analysis

Three to five 5-week-old, NOD.LCMV and NOD.Ln82-LCMV female mice were i.v. injected with 0.5μg of gp33 or D^b^ binding control peptide (ASNENMETM). Two hours post-injection, whole thymus tissue was processed as described above for total RNA extraction. RNA samples were hybridized to Affymetrix Mouse Gene 1.0 ST Arrays (Affymetrix – Thermo Fisher Scientific). Average signal intensities for each probe set within arrays were calculated by and exported from Affymetrix’s Expression Console (Version 1.1) software using the RMA method. Two pairwise comparisons were used to statistically resolve gene expression differences between experimental groups using the R/maanova analysis package (32). Specifically, differentially expressed genes were detected by using F1, the classical F-statistic that only uses data from individual genes. Statistical significance levels of the pairwise comparisons were calculated by permutation analysis (1000 permutations) and adjusted for multiple testing using the false discovery rate (FDR), q-value, method (33). Differentially expressed genes are declared at an FDR q-value threshold of 0.05. Two contrast groups were established: control D^b^ peptide (NOD.LCMV vs NOD.Ln82-LCMV), and gp33 peptide (NOD.LCMV vs NOD.Ln82-LCMV). Data were filtered for an expression fold change (EFC) > 2 and an expression FDR (q-value) < 0.05. Specific genes differentially regulated between NOD.LCMV and NOD.Ln82-LCMV are documented in Supplementary Table 1.

### Gene Expression analysis of NF-κB family members and NF-κB regulated genes

Live DP thymocytes were sorted from 5-6-week-old NOD-AI4 and NOD-AI4-*Nfkbid^−/−^*mice directly into fetal bovine serum using a FACSAria II sorter (BD Biosciences San Jose, CA, USA). Cells were pelleted, resuspended in TRIzol Reagent (Thermo Fisher Scientific) and frozen until use. Following phenol-chloroform phase separation, RNA was purified from the aqueous phase using a Quick-RNA MiniPrep Plus column (Zymo Research, Irvine, CA) according to the manufacturer’s protocol. Following RNA quantitation and reverse transcription as described above, qPCR was conducted using Power SYBR Green PCR master mix (Applied Biosystems – Thermo Fisher Scientific) with 250nM primers and 6ng of cDNA template. Primers are documented in Supplementary Table 2. Standard curves for each gene were performed using pooled cDNA, gene expression was normalized to *Gapdh* and expressed as fold change compared to NOD-AI4 control samples.

### Flow cytometry analysis

Single cell suspensions of thymus (Thy), spleen (SPL) and pancreatic lymph nodes (PLN) from 5-6-week-old female mice were prepared. Red blood cells in splenocyte samples were lysed with Gey’s buffer (34). Aliquots of 2×10^6^ cells were stained with the following monoclonal antibodies: CD4-BV650 (GK1.5) or -FITC (GK1.5), CD8-PE-Cy7 (53-6.72) or - APC (53-6.72), TCRVα8.3-FITC or -PE (B21.14), TCRVβ2-A647 (B20.6), TCRVβ8.1,2,3-FITC (F23.1) CD19-A700 (ID3), CD11b-PB (M1/70), CD11c-PB (N418), CD21-FITC (7G6), CD23-PE (B3B4), CD5-PE (53-7.313) acquired from BD Bioscience or BioLegend (San Diego, CA, USA), MHC class I tetramers Tet-AI4-PE (H2-D^b^/MimA2, sequence YAIENYLEL) (35) and Tet-NRPV7-PE (H2-K^d^/NRPV7, sequence KYNKANVFL) acquired from the National Institutes of Health tetramer core facility (Atlanta, GA, USA). Dead cells were excluded by DAPI or propidium iodide (PI) staining. Stained cells were acquired using a FACS LSRII instrument (BD Bioscience). All flow cytometric data were analyzed with FlowJo (FlowJo LLC, Ashland, OR, USA).

### T-cell Proliferation assay

Total splenic T-cells were purified from 5-week-old NOD and NOD-*Nfkbid*^−/−^ female mice by negative depletion of CD11c^+^, CD11b^+^, B220^+^, Ter19^+^, CD49b^+^ cells with biotin-conjugated antibodies and streptavidin magnetic beads over MACS columns (Miltenyi Biotech, Cologne, Germany) following manufacturer’s protocols. Purified T-cells were washed with cold PBS, counted and adjusted to a concentration of 2×10^7^/ml for labeling with 10μM cell tracer eFluor670 (eBioscience - Thermo Fisher Scientific). Labeled cells were adjusted to 4×10^6^/ml. Whole collagenase digested splenocytes from NOD.*scid* mice were used as APCs. Labeled T-cells were seeded in triplicate into a 96 well plate at a density of 2×10^5^/well with 4×10^5^ APC/well. Anti-CD3 monoclonal antibody (145-2C11) was added at final concentrations of 4, 2, 1, 0.5, 0.25 and 0.125μg/ml. The cells were incubated at 37°C, 5% CO_2_ for 48h. T-cell proliferation was assessed by flow cytometric analysis of cell tracer eFluor670 dilution. Proliferation parameters were obtained using the proliferation platform analysis in FlowJo (FlowJo LLC).

### Cytokine secretion analysis

IL-2 and IFN-γ secretion was assessed in supernatants of T-cell proliferation cultures by ELISA using BD OptEIA kits (BD Bioscience).

### Western blot

Whole thymus lysates were prepared in RIPA buffer (Cell Signaling Technologies, Danvers, MA, USA) containing Halt^TM^ Protease Inhibitor Cocktail (Thermo Fisher Scientific). Total protein content was quantified by Bradford assay (Thermo Fisher Scientific) and adjusted to 4mg/ml in RIPA buffer. Protein lysates were prepared for automated western blot using the Simple Wes system (ProteinSimple, San Jose, CA, USA). Briefly, samples were mixed at a 4:1 ratio with 5x fluorescent master mix and heated at 95°C for 5□min. Samples plus biotinylated molecular weight standards (ProteinSimple) were loaded along with blocking solution, biotin labeling reagent, wash buffers, primary antibodies, horseradish-peroxidase conjugated secondary antibodies and chemiluminescent substrate into a plate prefilled with stacking and separation matrices. In order to perform total protein normalization, each sample was loaded in duplicates; one capillary was used for immunodetection of Nfkbid and the second for total protein quantification. A 1:100 dilution of rabbit polyclonal anti-Nfkbid (36) with a 60 min incubation time was used for immunodetection. Chemiluminescent signal was captured and the resulting image was analyzed by the Compass for Simple Western software package (ProteinSimple). Nfkbid peak area was normalized to total protein area following ProteinSimple recommendations and standard guidelines (37, 38). Specifically the Nfkbid area for each sample was multiplied by a normalization factor (calculated as the total protein for the B6 sample divided by the total protein for each individual sample).

### Diabetes incidence

Diabetes was monitored once weekly using urine glucose strips (Diastix, Bayer, Leverkusen, Germany). Mice with two consecutive readings >250mg/dl (corresponding to a blood glucose of >300mg/dl) were considered diabetic.

### Statistical analysis

Data analysis and graphs were made using Prism 6 software (GraphPad, San Diego, CA, USA). Statistical analyses are detailed in the corresponding Figure Legends.

## Results

### Mapping an Idd7 region gene(s) regulating thymic numbers of diabetogenic AI4 CD8^+^ T-cells to a 5.4 Mb region on Chr. 7

We previously developed a mouse stock, designated NOD.*Chr7^B6^*FL-AI4, expressing the transgenic AI4 TCR and congenic for a B6-derived segment of Chr.7 overlapping the previously identified *Idd7* locus delineated by the flanking markers *Gpi1^b^*(34.2 Mb) and *D7Mit346* (58.7Mb) (29). AI4 T-cells underwent significantly higher levels of thymic deletion at the DP stage of development in the NOD.*Chr7^B6^*FL-AI4 congenic stock than in full NOD background mice (8). This indicated a gene(s) allelically varying between NOD and B6 mice within the overlapping congenic interval of the *Idd7* locus played a strong role in controlling the extent that diabetogenic AI4 CD8^+^ T-cells underwent thymic deletion (8). We subsequently derived from the original NOD.*Chr7^B6^*FL-AI4 stock two sub-congenic lines designated NOD.Ln82-AI4 (markers *D7Mit117*–*D7Mit247*) and NOD.Ln16-AI4 (markers *D7Mit79*–*D7Mit247*) (Fig. 1A). Flow cytometric analyses found that compared to NOD-AI4 controls, the yield of DP AI4 thymocytes (identified by staining with the TCR Vα8.3 specific monoclonal antibody B21.14) (Fig. 1B) was significantly decreased in the NOD.Ln82-AI4 sub-strain (Fig. 1C). In contrast, the yield of AI4 DP thymocytes in the NOD.Ln16-AI4 stock were similar to that in NOD-AI4 controls (Fig. 1B, C). These differences indicated a polymorphic gene(s) influencing the extent to which DP AI4 thymocytes undergo negative selection resides within a 5.4 Mb region on Chr.7 delineated by the markers *D7Mit117*-*D7Mit225*. Expression levels of the clonotypic Vα8.3 TCR element was higher in the NOD.Ln82-AI4 stock compared to NOD-AI4 controls (Fig. 1D), a trait previously observed in NOD.*Chr7^B6^*FL-AI4 mice (8). These results indicated a gene(s) in the 5.4 Mb region on Chr. 7 between *D7Mit117* and *D7Mit225* may modulate how efficiently AI4 T-cells undergo thymic deletion through regulation of TCR expression levels (8).

**Figure 1:**
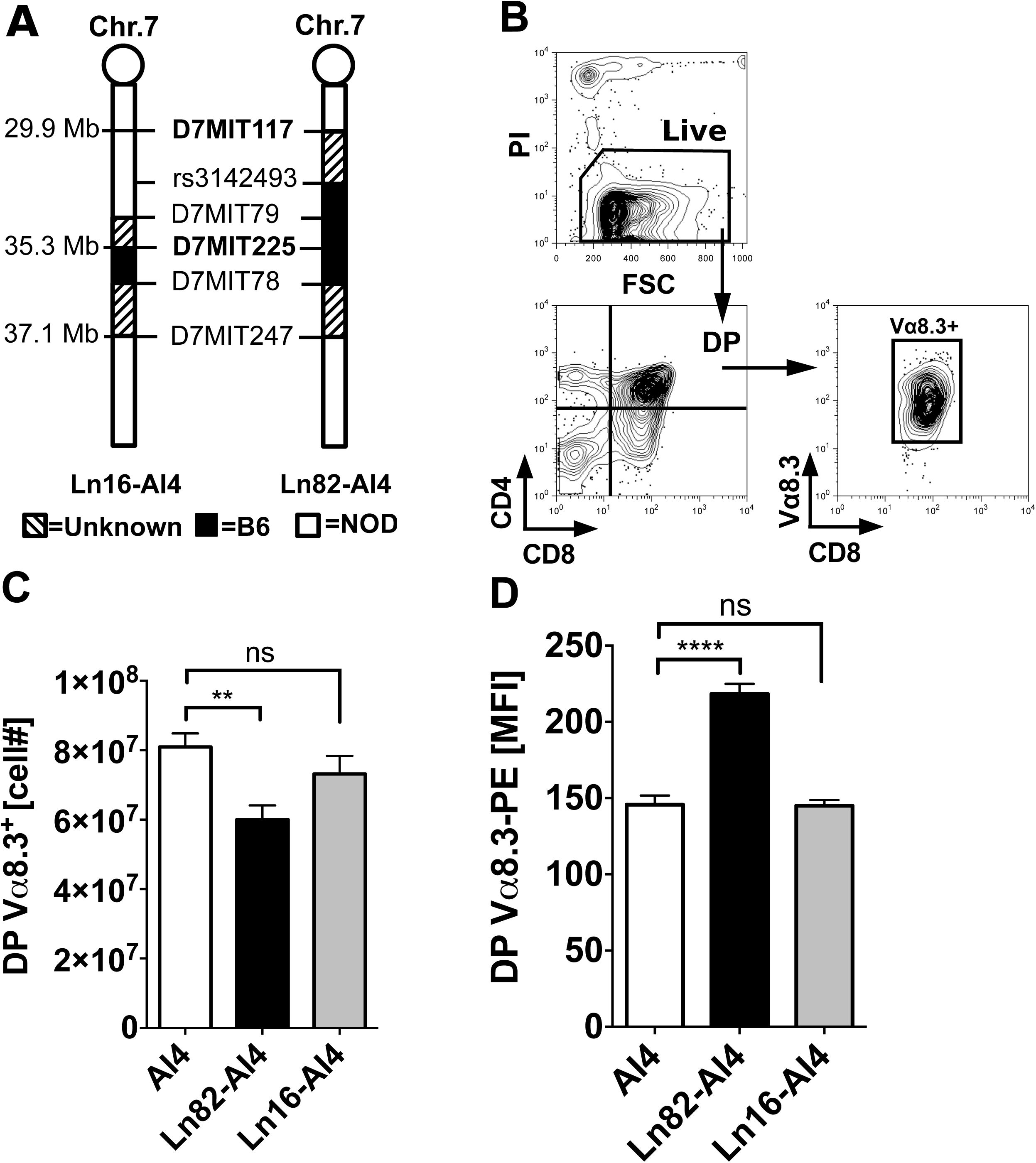
Mapping an *Idd7* region gene(s) controlling the thymic negative selection efficiency of diabetogenic AI4 CD8^+^ T-cells to a 5.4 Mb region on Chromosome 7. (**A**) Schematic diagram of B6 derived Chr.7 congenic regions in NOD-AI4 mouse strains. Marker positions are indicated in Mb based on genome assembly release GRCm38.p5 (40) (**B**) Flow cytometric gating strategy to enumerate AI4 DP thymocytes. Thymocytes from 5-week-old female NOD-AI4 (n=32), NOD.Ln82-AI4 (n=20) and NOD.Ln16-AI4 (n=14) were stained with CD4, CD8 and Vα8.3 specific antibodies. (**C**) Numbers of live (PI negative) CD4^+^CD8^+^ (DP) AI4 (Vα8.3^+^) thymocytes in the indicated strains. (**D**) Mean fluorescence intensity (MFI) of staining of DP thymocytes from the indicated strains by the monoclonal antibody Vα8.3-PE. All values represent mean±SEM of more than 3 independent experiments, statistical significance was determined by Log-rank (Mantel-Cox) t-test (ns p > 0.05, ** p < 0.001, **** p < 0.00001).

### Nfkbid is the only differentially expressed Idd7 region gene in thymocytes undergoing low versus high levels of deletion

Due to differing cellular composition, we did not feel it appropriate to utilize thymii from NOD-AI4 and NOD.Ln82-AI4 mice to carry out initial mRNA transcript analyses to identify genes potentially contributing to the differential negative selection efficiency of diabetogenic CD8^+^ T-cells in these strains. Instead, we reasoned a more appropriate platform to carry out such analyses would be provided by a previously produced NOD background stock transgenically expressing a Vα2^+^Vβ8^+^ TCR recognizing the H2-D^b^ class I restricted gp33 peptide (KAVYNFATM) derived from Lymphocytic Choriomeningitis Virus (LCMV) (26, 27). The basis for choosing this NOD.LCMV stock was that LCMV reactive CD8^+^ T-cells developing in the thymus would not be under any thymic negative selection pressure until such time their cognate antigen is exogenously introduced. We generated an additional NOD.LCMV stock homozygous for the Ln82 congenic interval. NOD.LCMV and NOD.Ln82-LCMV mice were i.v. injected with 0.5µg of the gp33 or an H2-D^b^ binding control peptide (ASNENMETM). At 24 hours post-injection, we assessed the proportion of Vα2^+^ TCR LCMV-specific DP thymocytes present in NOD.LCMV and NOD.Ln82-LCMV mice that had received the gp33 peptide relative to control cohorts. A significantly greater proportion of gp33 reactive DP thymocytes were deleted in NOD.Ln82-LCMV than NOD.LCMV mice (Fig. 2A). These results further indicated a polymorphic gene(s) in the *Idd7* region regulates the sensitivity of DP thymocytes to undergo antigen-induced deletion.

**Figure 2:**
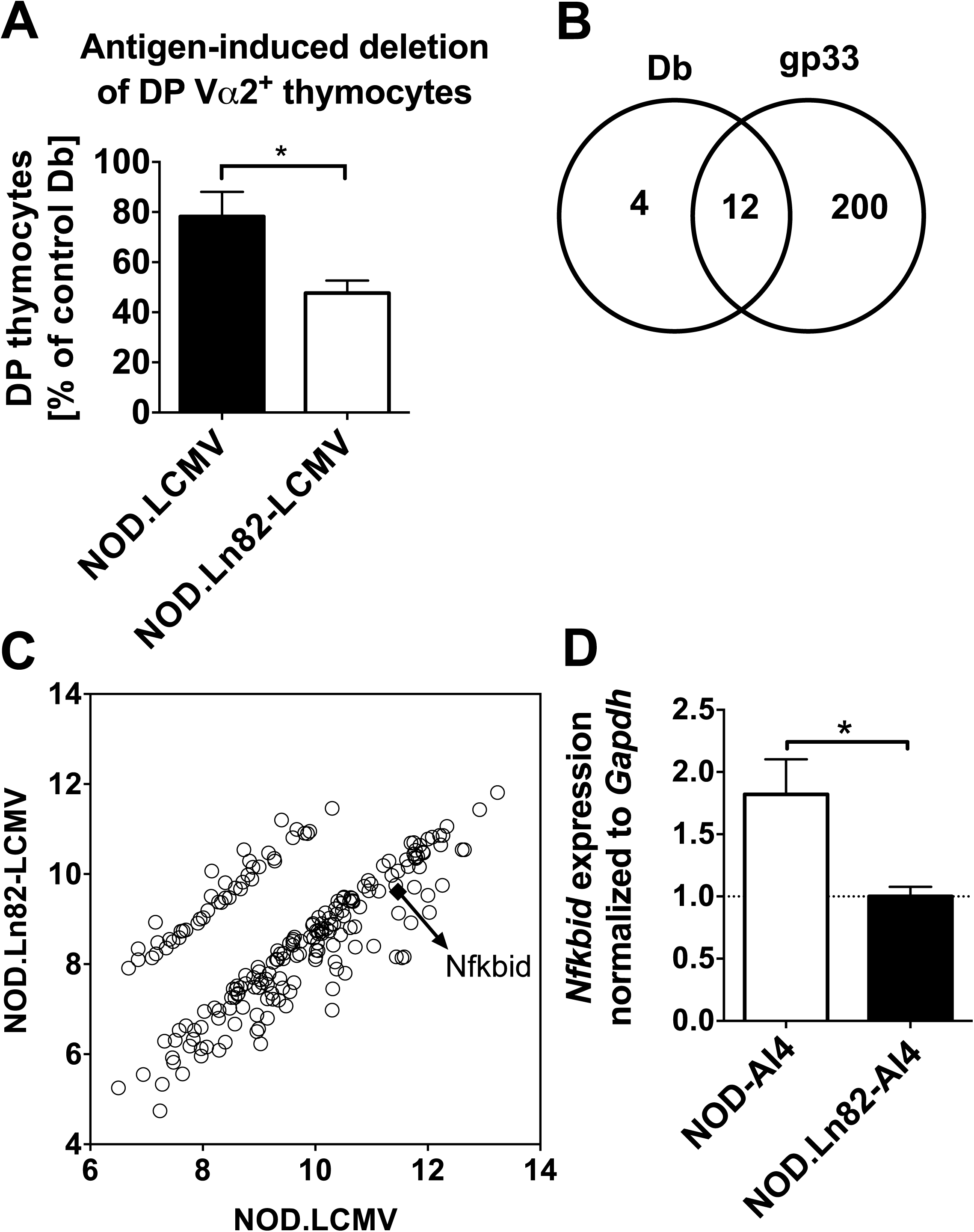
*Nfkbid* is the only differentially expressed *Idd7* region gene in diabetogenic CD8^+^ T-cells undergoing low versus high levels of thymic deletion. (**A**) For antigen-induced negative selection, either 0.5 μg of gp33 (cognate antigen) or D^b^ binding control peptide were i.v. injected into five-week-old female NOD.LCMV (n=18 for controls and n=6 for gp33 treated) or NOD.Ln82-LCMV (n=22 for controls and n=6 for gp33 treated) mice. Remaining DP Vα2^+^ thymocytes were quantified at 24 hours post-injection in each group. The efficiency of deletion was calculated as a ratio of the number DP Vα2^+^ thymocytes remaining in gp33 treated mice to the average numbers in controls and expressed as a percentage 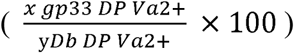. Statistical significance was determined by Log-rank (Mantel-cox) t-test (ns p > 0.05, * p < 0.05). (**B**) As assessed by microarray analyses number of genes deferentially expressed declared at a false discovery rate (q-value) <0.05 and a fold change ≥2 in thymocytes from NOD.LCMV and NOD.Ln82-LCMV 2 hours after i.v. injection with Db control or gp33 peptide. (**C**) Graph showing the average log_2_ (Intensity of probe set) for differentially expressed probe sets (same criteria as noted for panel B). The arrow identifies *Nfkbid*. Other genes are identified in Supplementary Table 1. (**D**) Total RNA was extracted from 2×10^5^ DP Vß8^+^ thymocytes sorted from 6-week-old female NOD-AI4 and NOD.Ln82-AI4 mice. *Nfkbid* mRNA relative quantification was done using the ddCt method using *Gapdh* as an endogenous control and NOD.Ln82-AI4 mice as reference, mean relative fold change (RFC) ± SEM is indicated. * p <0.05 unpaired t-test.

The LCMV TCR transgenic system provided an opportunity to compare the influence of the *Idd7* region over the transcriptional profile of thymocytes undergoing different levels of antigen-induced negative selection. Time-course experiments indicated that at 2 and 4 hours post-injection with the cognate antigen or peptide control there was no significant reduction in the number of Vα2^+^ DP thymocytes (Supplementary Fig. 2). However, we hypothesized that at 2-hours post-antigen injection, a differential gene expression profile may have already been induced in NOD.LCMV and NOD.Ln82-LCMV thymocytes contributing to their subsequent low and high level of deletion seen at 24 hours. Therefore, we conducted microarray analysis of whole thymus comparing NOD.LCMV and NOD.Ln82-LCMV at this time point. Using cutoff criteria of a q-value <0.05, and minimum two-fold strain variation, we found 16 and 212 genes were differentially expressed in thymocytes from NOD.LCMV and NOD.Ln82-LCMV mice injected with the D^b^ binding control or gp33 peptide respectively (Fig. 2B, Supplementary Table 1). This included 12 genes that were commonly differentially expressed in thymocytes from NOD.LCMV and NOD.Ln82-LCMV mice treated with either the control or gp33 peptide (Fig. 2B, Supplementary Table 1). None of the genes differentially expressed in thymocytes from NOD.LCMV and NOD.Ln82-LCMV mice treated with the control peptide mapped to the earlier defined 5.4 Mb *Idd7* support interval (genomic coordinates 7:29,926,306-35,345,930). Intriguingly, *Nfkbid* was the only differentially expressed gene mapping to this 5.4 Mb *Idd7* support interval in thymocytes from NOD.LCMV and NOD.Ln82-LCMV mice treated with the gp33 peptide (Fig. 3C). *Nfkbid* was expressed at a 3.6-fold higher level in thymocytes from gp33 treated NOD.LCMV than NOD.Ln82-LCMV mice (Fig. 3C). We also subsequently found by qPCR analyses that *Nfkbid* is expressed at ∼2-fold higher levels in flow cytometrically sorted equalized numbers of DP thymocytes from NOD-AI4 than NOD.Ln82-AI4 mice (Fig. 2D).

**Figure 3:**
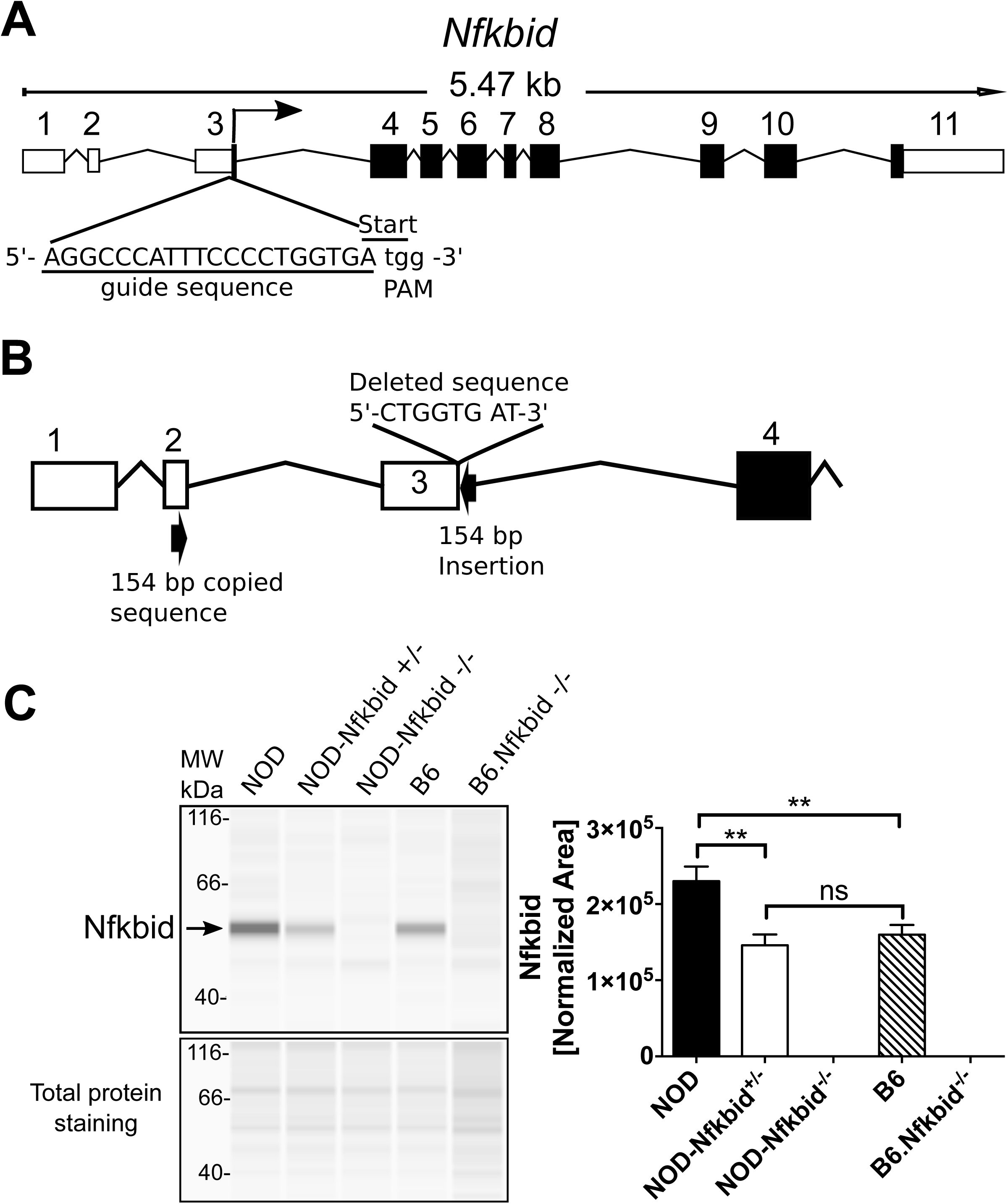
Targeting *Nfkbid* by CRISPR/Cas9 in NOD mice. (**A**) Representation of mouse *Nfkbid* gene showing the translation start site at the end of exon 3 and the sgRNA (underlined) designed to target this region by CRISPR/Cas9. The PAM sequence is indicated in lower case. (**B**) Schematic of the resulting targeted mutation consisting of an 8-nucleotide deletion and an insertion of 154 nucleotides originated by copy and inversion of a segment from *Nfkbid* exon 2 (For more detail, see Supplementary Figure 1). (**C**) Nfkbid protein abundance was assessed in total thymus lysates from the indicated strains by Simple Wes automated western blot analysis. B6.*Nfkbid*^−/−^ thymus samples were provided by Dr. Ingo Schmitz. Nfkbid was detected as a 55 kDa band. The difference in molecular weight compared to the theoretical 35 kDa, could be due to posttranslational modifications. Lower panel indicates the corresponding total protein staining for each sample. Right panel shows the Nfkbid-normalized area for NOD (n=6), NOD-*Nfkbid*^+/−^ (n=9), NOD-*Nfkbid*^−/−^ (n=8), B6 (n=6), and B6.*Nfkbid*^−/−^ (n=4) thymus samples. Nfkbid-peak area was normalized as follows: 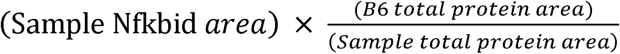. Bars represent the mean ± SEM. ns p >0.05, ** p <0.01, Mann-Whitney analysis.

*Nfkbid* (aka *I*κ*BNS*) encodes a non-conventional modulator of the transcription factor NF-κB (20) and ironically, was first discovered as a gene expressed in thymocytes undergoing negative, but not positive selection (21). Depending on cell type and physiological context, Nfkbid can stimulate or inhibit NF-κB pathway activity (20). Low to moderate levels of NF-κB activity (primarily the p50/p65 heterodimeric form) reportedly inhibits thymic negative selection of CD8^+^ lineage-destined T-cells (39). On the other hand, high NF-κB activity levels can elicit thymic negative selection of CD8^+^ T-cells (39). Thus, Nfkbid-mediated modulation of NF-κB activity could potentially contribute to the extent of diabetogenic CD8^+^ T-cell thymic deletion. In our microarray analysis, we found that 51 direct NF-κB-target genes, described in Dr. Thomas Gilmore’s curated database (http://www.bu.edu/nf-kb/gene-resources/target-genes/, accessed 3/26/2018), were upregulated versus none that were repressed in NOD.LCMV compared to congenic NOD.Ln82-LCMV thymocytes upon antigen stimulation (Supplementary Table 1). This indicated higher *Nfkbid* levels may be critical for upregulating NF-κB-mediated gene expression in antigen stimulated thymocytes. For this reason, we hypothesized that the higher *Nfkbid* expression levels characterizing antigen stimulated NOD thymocytes may elicit up-regulation of NF-κB target genes rendering these cells less prone to negative selection than those from B6 mice.

### Generation of NOD-Nfkbid deficient mice

In order to test whether the higher *Nfkbid* expression in NOD mice contributes to diminished thymic deletion of diabetogenic CD8^+^ T-cells, we utilized a CRISPR/Cas9 approach to ablate this gene. The mouse *Nfkbid* gene encodes four mRNA transcripts, three of which generate the same 327 amino acid protein, with a fourth representing a non-coding transcript (40). Based on the reference sequence (NM_172142.3), *Nfkbid* consists of 11 exons, with the start codon located at the end of exon 3 (Fig. 3A). We utilized a single guide RNA (sgRNA) (Fig. 3A) designed to disrupt the start codon in exon 3 and thereby eliminate translation of any potential transcripts. A resultant mutation consisting of an 8-nucleotide deletion coupled with an inverted insertion of a 154 nucleotide sequence copied from exon 2 was identified (Fig. 3B, supplemental Fig. 1) and subsequently fixed to homozygosity in a new stock officially designated NOD/ShiLtDvs-*Nfkbid^<EM3DVS>^*/Dvs and for simplicity abbreviated hereafter as NOD-*Nfkbid*^−/−^. Western blot analyses using total thymus lysates indicated Nfkbid was clearly present in NOD controls while being completely absent in both NOD-*Nfkbid*^−/−^ and previously generated (22, 31) B6.*Nfkbid*^−/−^ mice (Fig. 3C). Quantitative western blot analyses correlating with the earlier described microarray studies indicated thymic Nfkbid levels are significantly higher in NOD than B6 mice (Fig. 3C, right panel). Of potential physiological significance thymic Nfkbid protein levels in heterozygous NOD-*Nfkbid*^+/−^ mice were similar to that of the B6 strain. We did find transcription from the targeted allele is possible but does not produce Nfkbid protein, as shown in homozygous mutants (Supplementary Figure 1C, Figure 3C).

### Diminution of Nfkbid expression decreases thymic mRNA transcript levels of both some NF-κB subunit and target genes associated with enhanced negative selection of AI4 and NY8.3 diabetogenic CD8^+^ T-cells in NOD mice

We next evaluated whether diminution of *Nfkbid* expression enhanced the thymic deletion of diabetogenic CD8^+^ T-cells in NOD mice. To initially do this, the inactivated *Nfkbid* gene was transferred in both a heterozygous and homozygous state to the NOD-AI4 strain. Quantitation analyses of *Nfkbid* mRNA transcripts utilized primers that would only detect expression of the wild type allele. As expected, analysis of *Nfkbid* gene expression by qPCR showed respective high and absent *Nfkbid* wild type mRNA transcript levels in thymocytes from NOD-AI4 and NOD-AI4-*Nfkbid^−/−^* mice (Fig. 4A). Also, as expected, wild type *Nfkbid* mRNA transcript levels were significantly higher in thymocytes from NOD-AI4 than B6 control mice (Fig. 4A). In agreement with analyses of protein levels, a finding of potential physiological relevance was that thymic wild type *Nfkbid* mRNA transcript expression in NOD-AI4-*Nfkbid^+/−^*heterozygous mice was reduced to that characterizing B6 controls (Fig. 4A). Numbers of DP, but not CD8^+^ SP thymocytes positively staining with the H2-D^b^/MimA2 tetramer capable of binding the AI4 TCR (Tet-AI4) (gating strategy shown in Fig. 4B) were significantly less in both heterozygous and homozygous *Nfkbid* knockout NOD mice, compared to gene-intact controls (Fig. 4C). The same result was obtained by staining with Vα8.3 and Vβ2 specific antibodies binding the AI4 TCRα- and β-chains (results not shown). As shown in Fig. 1D, TCR expression was significantly higher on DP thymocytes from NOD.*Chr7^B6^*Ln82-AI4 congenic mice than in controls in which the *Idd7* region was of NOD origin. In contrast, partial or complete ablation of *Nfkbid* did not alter AI4 TCR expression levels on either DP or CD8^+^ SP thymocytes (Fig. 4D). Thus, some *Idd7* region gene(s) other than *Nfkbid* regulates thymic TCR expression levels. These latter results also disprove an earlier hypothesis (8) that an *Idd7* region gene(s) may control the thymic deletion efficiency of diabetogenic CD8^+^ T-cell by modulating TCR expression levels.

**Figure 4:**
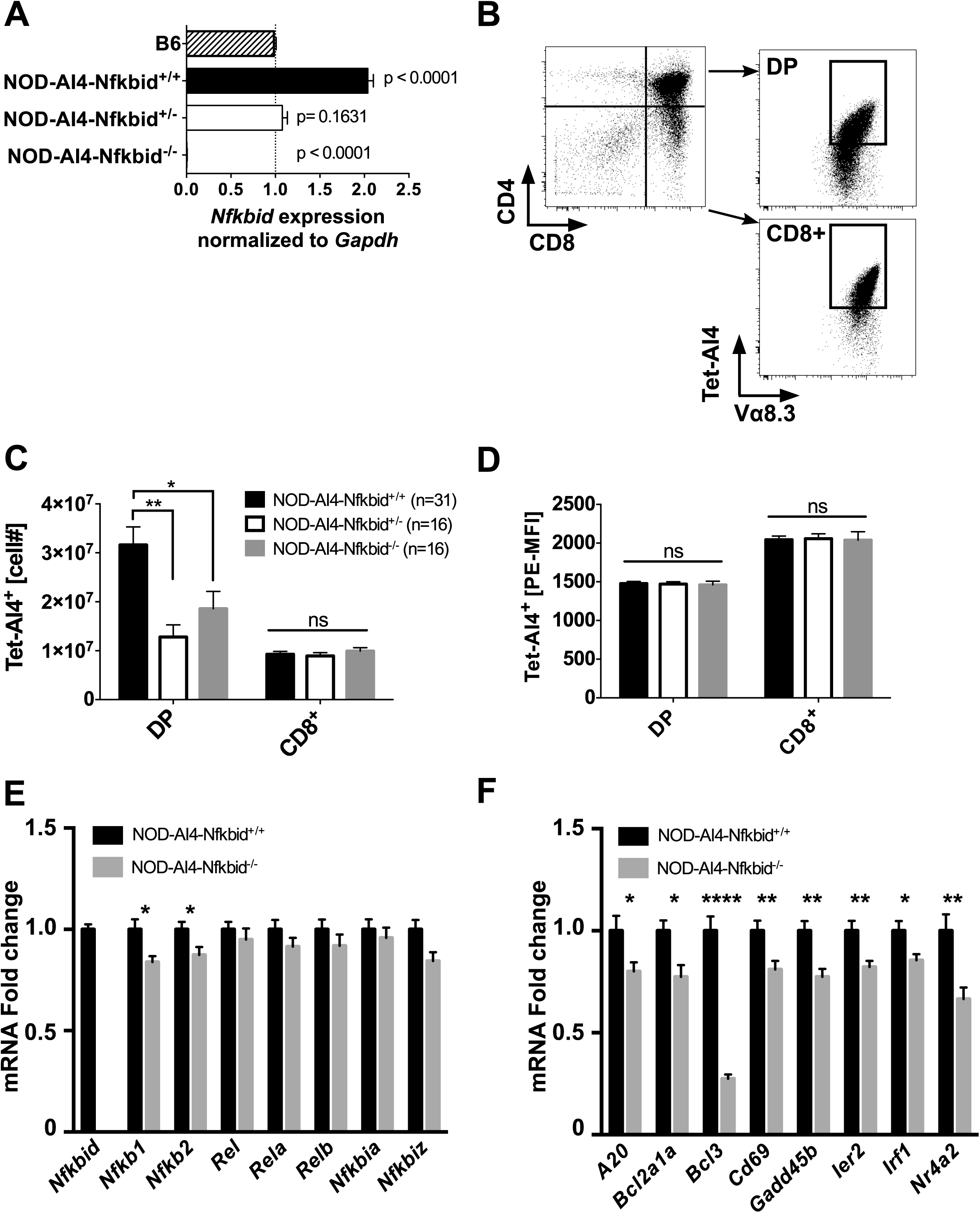
Genetic attenuation of *Nfkbid* expression controls the numbers of AI4 DP thymocytes. (**A**) Wild type *Nfkbid* gene expression was analyzed by qPCR of total thymus RNA from 6-week-old B6 (n=2), NOD-AI4 (n=4), NOD-AI4-*Nfkbid^+/−^* (n=4) and NOD-AI4-*Nfkbid^−/−^*(n=3) mice. *Gapdh* was used as endogenous control and B6 as the reference group. Graph shows the mean ± SEM of relative fold change of *Nfkbid* expression normalized to *Gapdh*. Statistical significance was determined by one-way ANOVA and Bonferroni’s multiple comparison test. The p-value of samples compared to the reference group is indicated. Note: We did not use B6.*H2^g7^*-AI4 mice as a reference in this study because the numbers of thymocytes is extremely low and would not provide a fair comparison for gene expression analysis. (B) Illustration of gating to enumerate DP and CD8^+^ SP thymocytes staining with the AI4 TCR specific tetramer. Depicted profiles are for thymocytes from an NOD-AI4 control. (C) Numbers of AI4 TCR expressing DP and CD8^+^ SP thymocytes from 6-week-old NOD-AI4, NOD-AI4-*Nfkbid^+/−^*, and NOD-AI4-*Nfkbid^−/−^* female mice. Data represent the mean ± SEM of DP or CD8^+^ SP thymocytes staining positive with Tet-AI4. (D) Geometric MFI of Tet-AI4 staining of the same DP and CD8**^+^** SP thymocytes evaluated in panel B. (E) Quantification of mRNA fold change expression of NF-κB family members in sorted DP NOD-AI4 vs NOD-AI4-*Nfkbid^−/−^* thymocytes. Individual gene expression was first normalized to *Gapdh*, and then fold change in comparison to expression in NOD-AI4 was calculated. Data is combined from 13 female mice, 5-6 weeks of age combined from two cohorts. (F) NF-κB target gene expression analysis was performed with the same samples as in (E). Bars represent the mean ± SEM. Statistical significance for A-D was analyzed by 2-way ANOVA and Bonferroni’s multiple comparison test. Statistical significance for E and F were analyzed by Mann-Whitney. ns p >0.05, * p <0.05, ** p <0.01, **** p<0.0001.

We next evaluated whether differential *Nfkbid* expression might regulate thymic negative selection efficiency by modulating the mRNA levels of NF-κB component members or downstream target genes we found they may control in the microarray study (Supplemental Table 1). We initially measured the gene expression of NF-κB family members in sorted DP thymocytes from NOD-AI4 and NOD-AI4-*Nfkbid*^−/−^ mice. As shown in Figure 4E, *Nfkb1* and *Nfkb2* gene expression levels were modestly decreased in *Nfkbid*^−/−^ mice, but other pathway members - *Rel*, *Rela* and *Relb* – were unchanged, as were the expression levels of two other nuclear inhibitors of NF-κB, *Nfkbia* and *Nfkbiz*. Additionally, we measured the expression levels of a subset of the 51 NF-κB target genes found to be upregulated to a greater extent in thymocytes from NOD.LCMV than NOD.Ln82-LCMV mice injected with gp33 peptide (Supplementary Table 1) to determine if they were also altered by ablation of *Nfkbid*. We found expression levels of a panel of NF-κB target genes were significantly lower in sorted DP thymocytes from NOD-AI4-*Nfkbid^−/−^* than NOD-AI4 mice (Fig. 4F). These results indicated Nfkbid likely acts as an enhancer of NF-κB pathway activity in this cellular context.

We next tested whether the apparent ability of varying *Nfkbid* expression levels to modulate thymic deletion efficiency of diabetogenic CD8^+^ T-cells extended to clonotypes beyond AI4. Thus, we introduced the *Nfkbid* knockout allele into a NOD stock with CD8^+^ T-cells transgenically expressing the NY8.3 TCR (NOD.NY8.3) that recognizes a peptide derived from the ß-cell protein islet-specific glucose-6-phosphatase catalytic subunit-related protein (IGRP) (28, 41). As observed for the AI4 clonotype, numbers of DP thymocytes staining with a tetramer (Tet-NRPV7) recognized by the NY8.3 TCR (Fig. 5A) were significantly less in 6-week-old NOD.NY8.3-*Nfkbid^+/−^*heterozygous mice than in controls (Fig. 5B). When examined at 4 weeks of age, numbers of both NY8.3 DP and CD8^+^ SP thymocytes were significantly less in NOD.NY8.3-*Nfkbid^−/−^* homozygous mice than in controls (Fig. 5C). Also, similar to the case observed for the AI4 clonotype, expression levels of the NY8.3 TCR on DP or CD8^+^ SP thymocytes was not altered by the presence or absence of Nfkbid (Fig. 5D). Thus, based on assessments of two separate clonotypes, we conclude that the *Nfkbid* expression variant characterizing NOD mice represents the *Idd7* region gene contributing to impaired thymic deletion of diabetogenic CD8^+^ T-cells in this strain.

**Figure 5:**
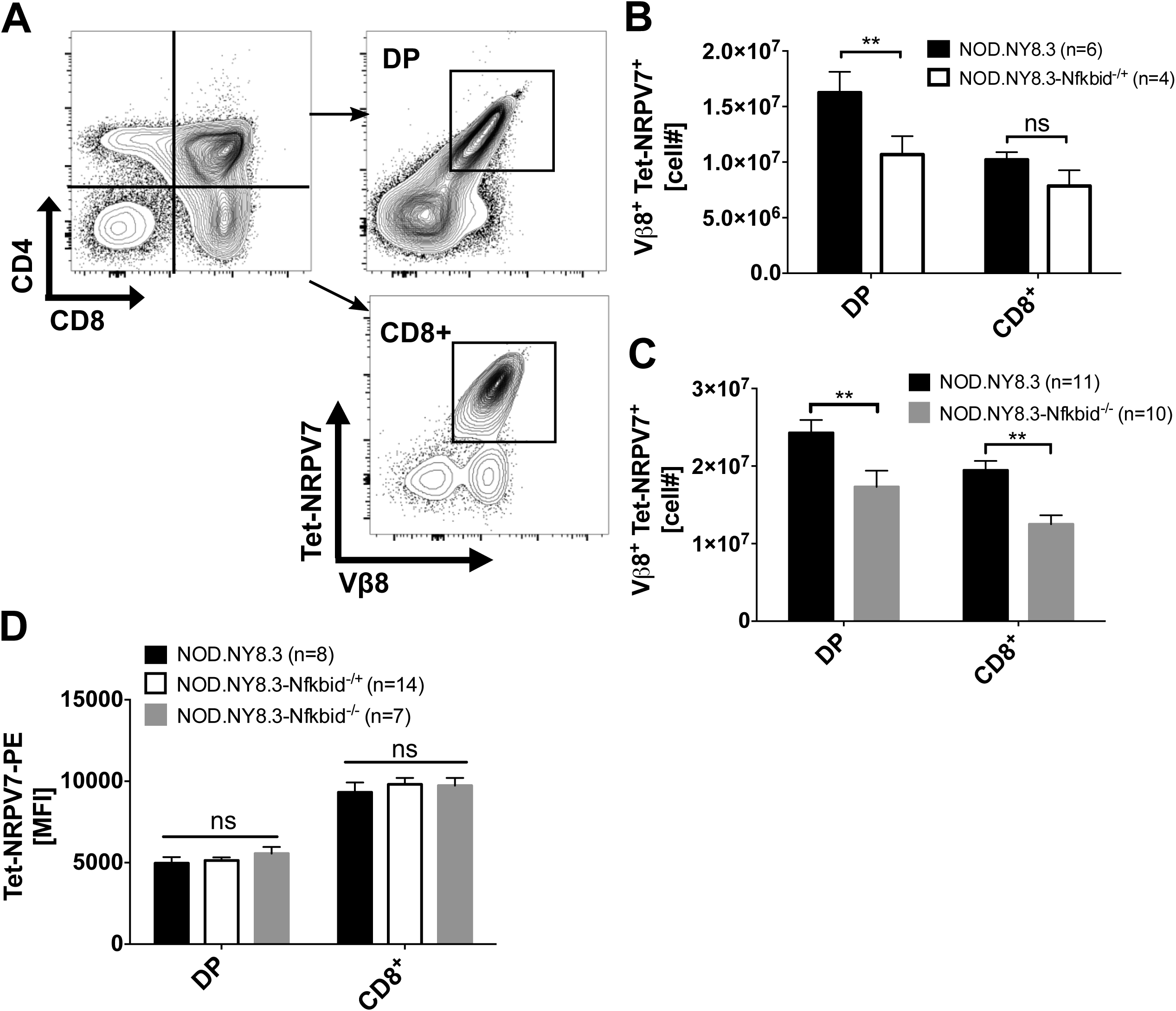
*Nfkbid* expression levels modulate thymic negative selection efficiency of diabetogenic NY8.3 clonotypic CD8^+^ T-cells. (A) Illustration of gating to enumerate DP and CD8^+^ thymocytes staining with the NY8.3 TCR specific Tet-NRPV7. Depicted profiles are for thymocytes from an NOD.NY8.3 control. (B) Numbers of NY8.3 TCR expressing DP and CD8^+^ thymocytes from 6-week-old female NOD.NY8.3 and NOD.NY8.3-*Nfkbid*^+/−^ heterozygous mice. Data represent the mean±SEM of DP or CD8**^+^** SP thymocytes staining positive with the NY8.3 specific Tet-NRPV7 reagent. (C) Numbers of NY8.3 TCR expressing DP and CD8**^+^** SP thymocytes from 4-week-old female NOD.NY8.3 and NOD.NY8.3-*Nfkbid*^−/−^ homozygous mice. Bars represent the mean±SEM of 3 independent experiments. (D) Geometric MFI of Tet-NRPV7 staining of DP and CD8**^+^** SP thymocytes from NOD.NY8.3, NOD.NY8.3-*Nfkbid*^+/−^, and NOD.NY8.3-*Nfkbid*^−/−^ mice. Bars represent mean±SEM. Statistical significance was analyzed by 2-way ANOVA and Bonferroni’s multiple comparison test. ns p >0.05, ** p <0.01.

### Nfkbid deficiency paradoxically contributes to accelerated T1D in NOD mice by impairing development of regulatory lymphocyte populations

We previously reported that *Idd7* region gene effects were restricted to the thymus. This was based on findings that while numerically reduced in the thymus, the AI4 T-cells that did reach the periphery of B6.*H2^g7^* and NOD.*Chr7^B6^*FL background mice were still able to expand and mediate T1D development (7, 8). However, we were surprised to find T1D development was actually accelerated in NOD-*Nfkbid^−/−^* mice (Fig. 6A). This prompted us to assess if ablation of the *Nfkbid* gene in NOD mice induced any numerical or functional differences in peripheral leukocyte populations varying from those previously reported in B6 mice carrying a similar mutation, but that exhibited no signs of autoimmunity (22, 31, 42). As previously observed for the B6 background stock, numbers of splenic CD4^+^ and CD8^+^ T-cells in NOD-*Nfkbid^−/−^* mice did not differ from wild type controls (data not shown). Also as previously observed in B6 (43), marginal zone (MZ) B-lymphocytes in the spleen and B1a cells in the peritoneum were both significantly reduced in *Nfkbid* deficient NOD mice (Fig. 6B, C). The proliferative capacity of CD4^+^ and CD8^+^ T-cells and their ability to secrete the cytokines IL-2 and IFNγ was reported to be diminished in *Nfkbid* deficient B6 mice (22). Similar phenotypes characterized CD4^+^ and CD8^+^ T-cells from NOD-*Nfkbid^−/−^* mice (Fig. 6D, E). Ablation of *Nfkbid* has also been reported to diminish numbers of FoxP3 expressing Tregs in B6 background mice (31). A similar effect was elicited by ablation of *Nfkbid* in the NOD strain (Fig. 6F). Previous reports indicated Nfkbid is a direct regulator of IL-10 (36). Proportions of FoxP3^+^ and FoxP3^−^ CD4^+^ T-cells producing immunosuppressive IL-10 upon *ex vivo* stimulation were significantly lower in *Nfkbid* deficient than intact NOD mice (Fig. 6G). We recently found that partially overlapping populations of CD73 expressing and IL-10 producing Bregs can inhibit autoimmune T1D development in NOD mice (44). Ablation of *Nfkbid* also reduced levels of such IL-10 producing and CD73 expressing Bregs in NOD mice (Fig. 6H, I). Thus, while diminishing *Nfkbid* expression enhances thymic negative selection of diabetogenic CD8^+^ T-cells in NOD mice, the coincidental decrease in numbers of immunoregulatory lymphocyte populations likely at least in part accounts for why any pathogenic effectors that do reach the periphery still aggressively expand and induce disease development.

**Figure 6:**
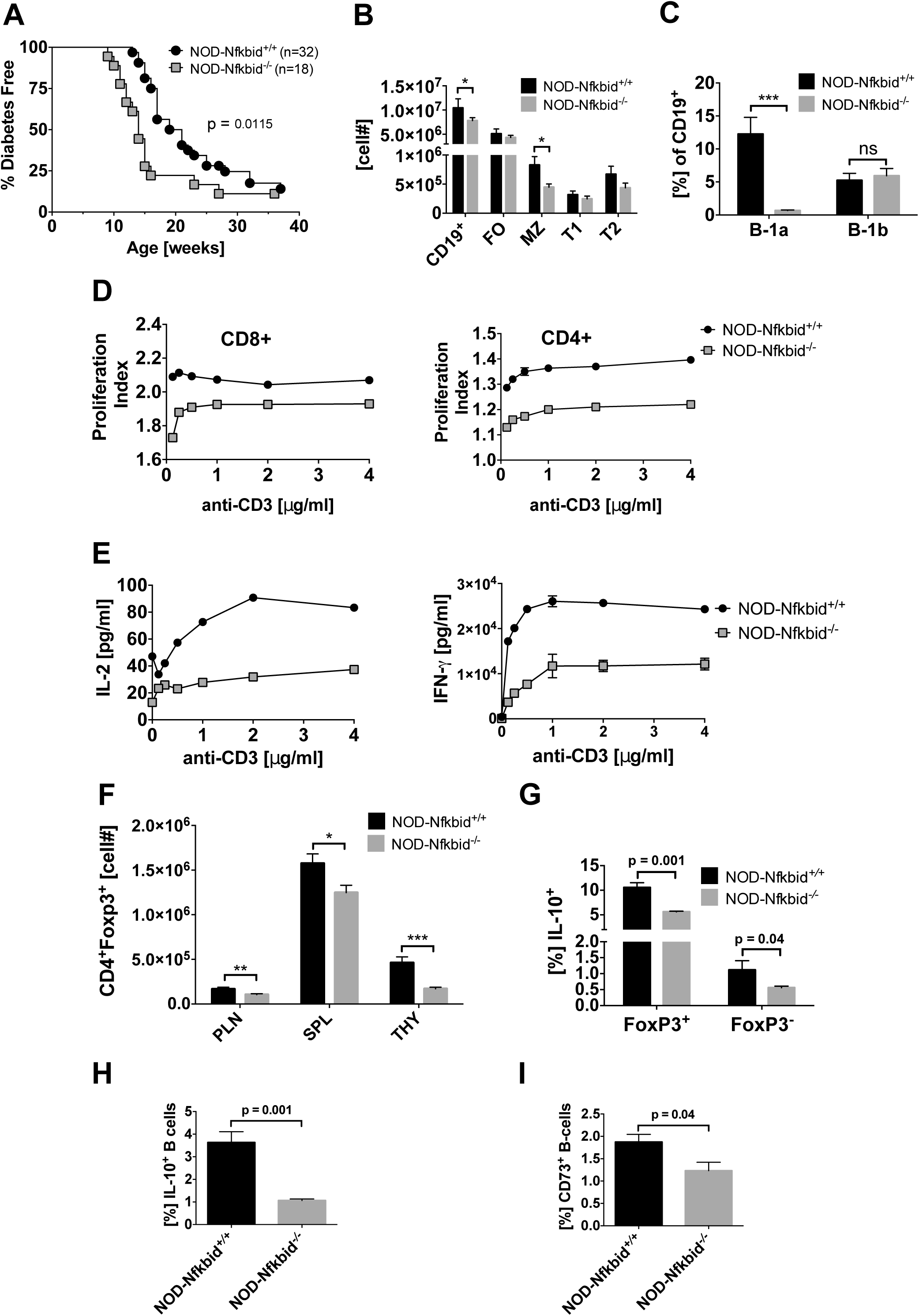
*Nfkbid* deficiency paradoxically accelerates T1D onset in NOD mice by impairing development of regulatory lymphocyte populations. (A) T1D development was analyzed in a cohort of NOD and NOD-*Nfkbid^−/−^* female mice. Survival curve comparisons were analyzed through Log-rank (Mantel-Cox) test. (B) Splenocytes from 7-week-old female NOD (n=3) and NOD-*Nfkbid^−/−^(*n=5) mice were labeled with antibody for CD19, CD21 and CD23 and analyzed by flow cytometry. Singlet, DAPI^−^ CD19^+^ cells were identified as total B-cells. Based on expression of CD23 and CD21, B-cells were classified as follicular (FO: CD23^+^CD21^lo^), marginal zone (MZ: CD23^−^CD21^hi^) transitional 1 (T1: CD23^−^ CD21^−^) and transitional 2 (T2: CD23^int^ CD21^hi^) (C) Peritoneal cavity B-cells were isolated by lavage with cold HBSS from the same group of mice described in (B), and stained for CD19, B220, CD5 and analyzed by flow cytometry. Total B-cells (Singlet, DAPI^−^ CD19^+^) were classified based in the expression of B220 and CD5 as B1-a (CD5^+^ B220^int/lo^) and B1-b (CD5^−^ B220^lo^) (D) The proliferation capacity of purified total T-cells from NOD and NOD-*Nfkbid^−/−^* mice was assessed as described in methods section. Briefly, MACS purified total T-cells were labeled with cell tracer eFluor670 and stimulated *in vitro* with increasing amounts of anti-CD3 monoclonal antibody. After 48 hours of culture the cells were labeled with antibody for CD4 and CD8. Proliferation in CD4^+^ and CD8^+^ T-cell subsets was analyzed by flow cytometric analyses of cell tracer eFluor670 dilution. Proliferation index was determined using the Flow Jo cell proliferation tool. Data represent the mean±SEM of 3 technical replicates. (E) Cell culture supernatants of purified T-cells used in the proliferation assay were assessed for secretion of the cytokines IL-2 and IFN-γ by ELISA. Data represent mean±SEM of 3 technical replicates. (F) Enumeration of CD4^+^ FoxP3^+^ (based on *Foxp3^tm2(eGFP)Tch^* reporter expression) Tregs in 6-week-old female NOD (n=5) and NOD-*Nfkbid^−/−^* mice (n=9) from pancreatic lymph node, spleen and thymus. (G) Analysis by intracellular staining and subsequent flow cytometric analyses of proportion of Foxp3^+^ and FoxP3^−^ splenic CD4^+^ T-cells from 6-week-old female NOD (n=5) and NOD-*Nfkbid^−/−^* mice (n=9) producing IL-10. (H) Analysis by intracellular staining and subsequent flow cytometric analyses of proportions of IL-10 producing splenic B-lymphocytes in 6-week-old female NOD (n=5) and NOD-*Nfkbid^−/−^* mice (n=9). (I) Proportions of splenic CD73^+^ B-lymphocytes in 6-week-old female NOD (n=5) and NOD-*Nfkbid^−/−^* mice (n=9). All bar graphs represent mean±SEM. Statistical significance was determined by 2-way ANOVA and Bonferroni’s multiple comparison test. ns p >0.05, * p <0.05, ** p <0.01, *** p <0.001.

## Discussion

Previous work from our group demonstrated the common class I variants expressed by NOD mice (H2-K^d^, H2-D^b^) have an aberrant lessened ability to mediate the thymic negative selection of diabetogenic CD8^+^ T-cells, primarily due to interactions with some non-MHC gene(s) in the *Idd7* locus on proximal Chr. 7 (7). In this study, through congenic truncation analyses, we fine mapped the *Idd7* region gene(s) controlling the efficiency of diabetogenic CD8^+^ T-cell thymic deletion to a 5.4 Mb interval on proximal Chr. 7. Identification of the responsible *Idd7* region gene was greatly aided by the exploitation of NOD.LCMV TCR transgenic stocks that did or did not carry a B6 derived congenic interval encompassing the relevant region of proximal Chr. 7. The advantage of these stocks is that their thymocytes are under no negative selection pressure until such time the LCMV cognate gp33 antigenic peptide is deliberately introduced. Upon introduction of the gp33 peptide, LCMV DP thymocytes were deleted to a significantly greater extent in the presence of B6- rather than NOD-derived genome across the relevant *Idd7* region. Subsequent microarray analyses found that following introduction of the gp33 antigenic peptide *Nfkbid* was the only differentially expressed gene mapping to the relevant 5.4 Mb region on Chr. 7. The presence of NOD rather than B6 derived genome across this *Idd7* region resulted in *Nfkbid* being expressed at significantly higher levels in antigen stimulated LCMV reactive thymocytes. This was similarly true for matched numbers of DP thymocytes transgenically expressing the TCR from the naturally occurring diabetogenic AI4 CD8^+^ T-cell clone derived from NOD mice. Thus, we used CRISPR/Cas9 technology to eliminate *Nfkbid* expression in NOD mice to see if improved thymic negative selection would result. NOD stocks in which Nfkbid protein was completely absent were found to have an improved capacity to mediate thymic negative selection of both TCR transgenic AI4 and NY8.3 CD8^+^ diabetogenic T-cells. Similar results were seen in heterozygous stocks where thymic Nfkbid protein levels were equivalent to that in B6 controls. These collective results indicated the higher *Nfkbid* expression variant in NOD than B6 mice is likely the *Idd7* region gene strongly contributing to diminished thymic deletion of diabetogenic CD8^+^ T-cells in the former strain.

Nfkbid is a mostly nuclear protein containing 7 ankyrin repeat domains originally reported as an inhibitor of NF-κB signaling (21). However, due to the lack of DNA binding and transactivation domains, Nfkbid function is ultimately mediated through protein-protein interactions with NF-κB family members. Accordingly, Nfkbid can function as an inhibitor or enhancer of NF-κB activities depending on the protein partner and cell type (20). This includes previously reported interactions of Nfkbid with the NF-κB subunit p50 (Nfkb1) in thymocytes undergoing negative selection (21). Our current results indicate *Nfkbid* is not required for thymocytes destined for the CD8^+^ lineage to undergo negative selection, and instead its particular expression level in NOD mice appears to diminish the efficiency of this process. Both our microarray analysis of NOD.LCMV thymocytes, as well as qPCR analysis of sorted NOD-AI4 DP thymocytes, indicate that in the context of negative selection of autoreactive CD8^+^ T-cells, Nfkbid likely acts as an enhancer of NF-κB activity. Nfkbid can be incorporated into NF-κB heterodimers with the p50, p52, or c-Rel binding partners (20). As we observed in this study with ablation of Nfkbid, an acceleration of T1D development concomitant with a decrease in Treg numbers has also been reported to characterize NOD mice made deficient in c-Rel expression (45). Thus, it is possible that in both cases T1D acceleration resulted from a decrease in c-Rel/Nfkbid heterodimer levels. The future dissection of such possible pathways in our NOD-AI4 model system is warranted to better understand how Nfkbid/NF-κB signaling contributes to CD8^+^ T-cell selection.

Three different *Nfkbid* deficient mouse models have been previously produced, all on the B6 background (22, 42, 46). These previously produced models have been useful to identify several functional aspects of *Nfkbid*. In the B-cell compartment, *Nfkbid* expression is necessary for development of the MZ and B1-a subsets as well as the generation of antibody responses to T-dependent and independent antigens (22, 43, 46–48). *Nfkbid* deficiency was also associated with an impairment in IL-10 secreting B-lymphocytes (49). *Nfkbid* deficient T-cells have impaired *in vitro* proliferation and cytokine secretion (IL-2, IFN-γ) (22). We found these phenotypes also characterize our *Nfkbid* deficient NOD mouse stock.

Interaction of Nfkbid with c-Rel has also been reported to contribute to Foxp3 expression in Treg cells (31). Thus, we found it of interest that in addition to enhancing the thymic deletion of diabetogenic CD8^+^ T-cells, diminution of *Nfkbid* expression also resulted in decreased numbers of peripheral Tregs in NOD mice. We recently reported that overlapping populations of CD73 expressing and IL-10 producing Bregs can exert T1D protective immunoregulatory activity in NOD mice (44). Ablation of *Nfkbid* expression also diminished levels of both CD73 expressing and IL-10 producing Bregs in the periphery. Thus, we conjecture it is the loss of both Tregs and Bregs that at least in part accounts for the unexpected finding that T1D development is actually accelerated in *Nfkbid* deficient NOD mice despite their improved, albeit not completely restored, ability to mediate the thymic deletion of pathogenic CD8^+^ T-cells. Hence, *Nfkbid* expression level appears to act as a two-edged sword, lower levels than that in the thymus of NOD mice can result in better negative selection of autoreactive CD8^+^ T-cells, but at the same time can contribute to autoimmunity in the periphery by diminishing levels of regulatory lymphocyte populations. Determining whether *Nfkbid* expression- or activity-levels can be fine-tuned to improve autoreactive CD8^+^ T-cell negative selection without diminishing Treg and Breg populations is warranted.

In conclusion, *Nfkbid* expression levels appear to be an important contributory factor to how efficiently autoreactive diabetogenic CD8^+^ T-cells undergo thymic negative selection. However, this study also revealed a caveat that would have to be considered if it might ultimately become possible to diminish *Nfkbid* expression through pharmacological means. That is while diminishing *Nfkbid* expression can allow for increased thymic deletion of diabetogenic CD8^+^ T-cells, it does so at the price of decreasing levels of regulatory lymphocytes allowing any pathogenic effectors that do reach the periphery to more rapidly induce disease onset. This explains our previous findings that the *Idd7* region exerts opposing effects where the presence of B6 derived genome in the region exerted T1D protective effects at the thymic level, but in the periphery actually promotes disease development (8). Thus, when contemplating possible future interventions designed to diminish numbers of diabetogenic T-cells, a consideration that should be taken into account is whether the treatment might also impair regulatory cell numbers or activity allowing the pathogenic effectors which do remain present to become more aggressive.

## Acknowledgements

We would like to thank the Jackson Laboratory Research Animal Facility, Genetic Engineering Technology group, Flow Cytometry service, and Transgenic Genotyping core for technical support on this project. We thank the National Institutes of Health tetramer core facility for the tetramers used in this study. The authors would also like to thank Carl Stiewe and his wife, Maike Rohde, for their generous donation toward T1D research at The Jackson Laboratory, which contributed to this work.

## Footnotes

1 MP was supported by JDRF Fellowship 3-PDF-2014-219-A-N. For parts of this work, JJR1 was supported by either NIH Fellowship 1F32DK111078 or JDRF Fellowship 3-PDF-2017-372-A-N. He was also supported by a grant from the Diabetes Research Connection DRC 006887 JR. DVS is supported by NIH grants DK-46266, DK-95735, and OD-020351-5022. This work was also partly supported by Cancer Center Support Grant CA34196. IS was supported by grants of the Deutsche Forschunggemeinschaft (SCHM1586/6-1 and project A23 of SFB854).

2 MP designed and conducted experimentation, interpreted data, and wrote the manuscript. JJR1, contributed to experimental design, data interpretation, and writing of the manuscript. JRD designed and conducted experiments, interpreted data and contributed to writing the manuscript. JJR2, DJL, JA and HDC conducted experimentation. VKS and TS contributed to statistical analyses. YGC, AG and IS contributed to experimental design. DVS contributed to study conception, supervised experimental effort, and writing of the manuscript.

3 C57BL/6J, (B6); CD4^+^CD8^+^ double positive, (DP); CD4^+^Foxp3^+^ regulatory T-cell, (Treg); chromosome, (Chr.); dendritic cell, (DC); Lymphocytic Choriomeningitis Virus, (LCMV); H2-D^b^/MimA2 tetramer capable of binding the AI4 TCR, (Tet-AI4); H2-K^d^/NRPV7 tetramer capable of binding the NY8.3 TCR, (Tet-NRPV7); islet-specific glucose-6-phosphatase catalytic subunit-related protein, (IGRP); NOD/ShiLtDvs, (NOD); NOD/ShiLtDvs-*Nfkbid*^<EM3DVS>/^Dvs, (NOD-*Nfkbid^−/−^);* NOD.129X1(Cg)-Foxp3^tm2Tch^/Dvs, (NOD.*Foxp3-eGFP)*; NOD.B6-(Gpi1-D7Mit346)/LtJ, (NOD.*Chr7^B6^*FL); NOD.Cg-*Prkdc^scid^*Emv30b/Dvs, (NOD.*scid*); pancreatic lymph node, (PLN); propidium iodide, (PI); regulatory B-cell, (Breg); single guide RNA, (sgRNA); type 1 diabetes, (T1D); spleen, (SPL); thymus, (Thy); type 1 diabetes susceptibility, (*Idd*).

