## Supplementary Materials for "A hypermorphic *Nfkbid* allele represents an *Idd7* locus gene contributing to impaired thymic deletion of autoreactive diabetogenic CD8^+^ T-cells in NOD mice"

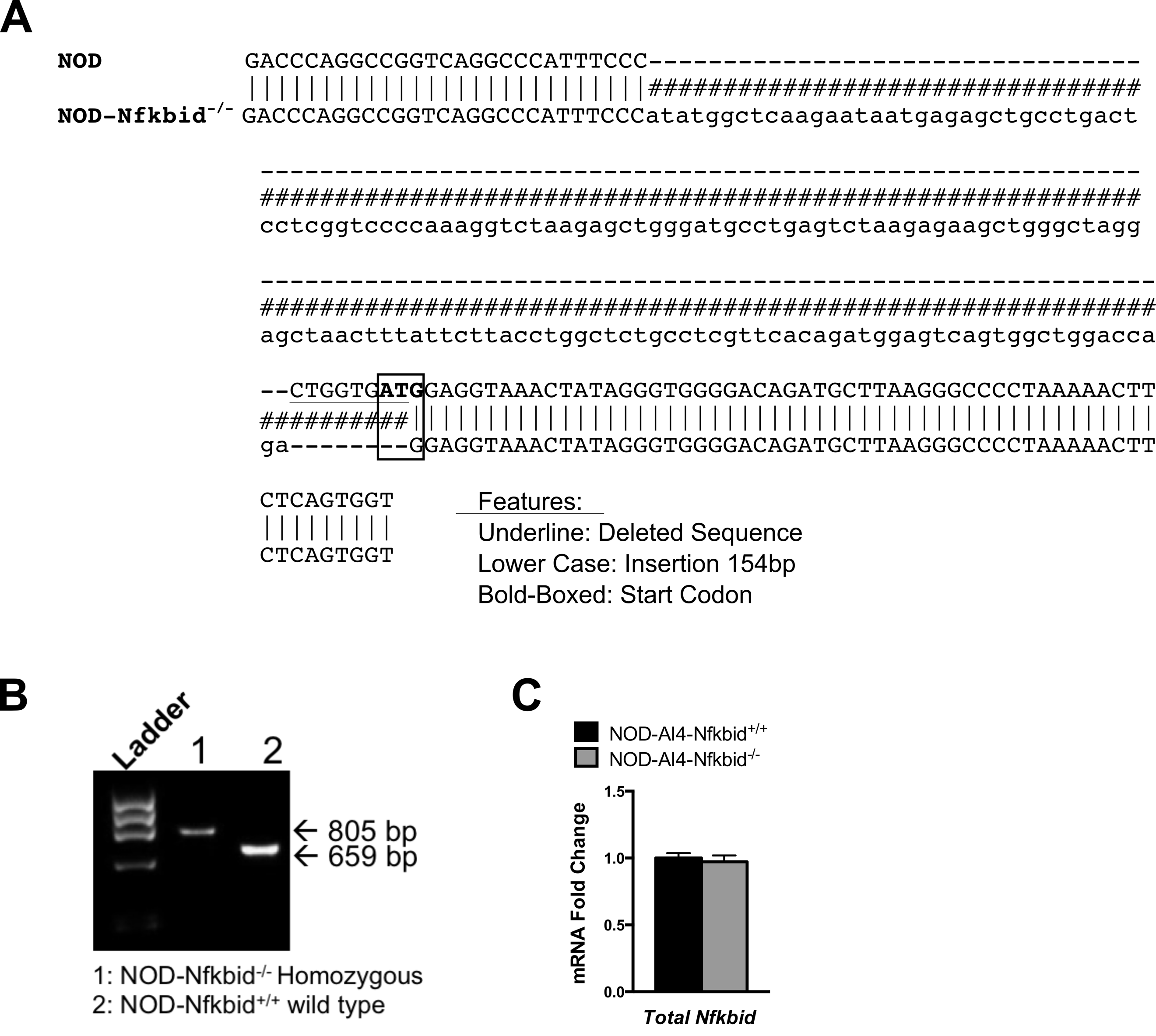

**Supplementary Figure 1 – *Nfkbid* knockout allele detail**

DNA from tail-tips was prepared and the region surrounding the start codon in exon 3 of *Nfkbid* in NOD or NOD-*Nfkbid^-/-^* mice was amplified (forward: 5’–GGAGACTCCAGGTTTAC–3’; reverse: 5’–CCAGGATGGGATCTTAAA–3’). Mutant amplicons were sequenced via Sanger sequencing and analyzed via gel electrophoresis. **(A)** Diagram showing alignment of the NOD-*Nfkbid^-/-^* mutant allele with the NOD *Nfkbid* sequence. Note the 8bp deletion (underlined) and 154bp insertion (lowercase) disrupting *Nfkbid*. **(B)** Representative 1.5% agarose gel showing 805bp mutant (Lane 1) amplicon compared to the 659bp wild type amplicon (Lane 2). **(C)** Total *Nfkbid* mRNA expression using a primer set that amplifies a region surrounding the junction of exon 6 and exon 7. Transcription from the mutant allele does not result in detectable Nfkbid protein (Figure 3C).

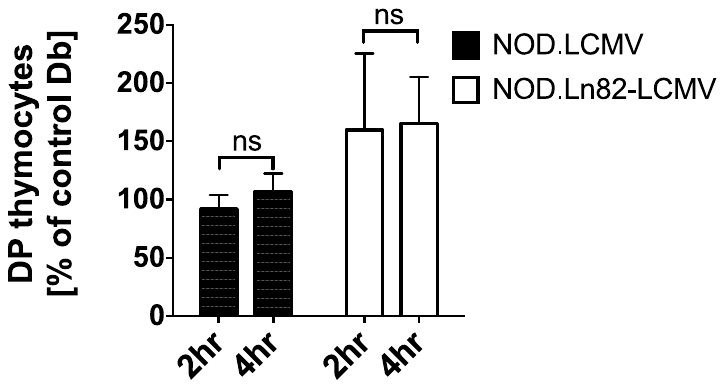

**Supplementary Figure 2 – Time course analysis of antigen-induced deletion of DP Vα2+ thymocytes upon injection with gp33 peptide.**

Five-week-old female NOD.LCMV or NOD.Ln82-LCMV were i.v. injected with 0.5μg of gp33 (cognate antigen) or Db binding control peptide (n=18 control, n=7 gp33 treated NOD.LCMV; n=22 control, n=6 gp33 treated NOD.Ln82-LCMV). Double positive (DP) Vα2+ thymocytes were quantified at 2 and 4 hours post injection for each group. The efficiency of deletion was calculated as a ratio of the number of DP Vα2+ thymocytes remaining in gp33 treated mice to the average numbers in controls (# gp33 DP Vα2+ / # Db DP Vα2+ x 100) and expressed as percentages. Statistical significance was determined by Log-rank (Mantel-Cox) t-test (ns p>0.05).

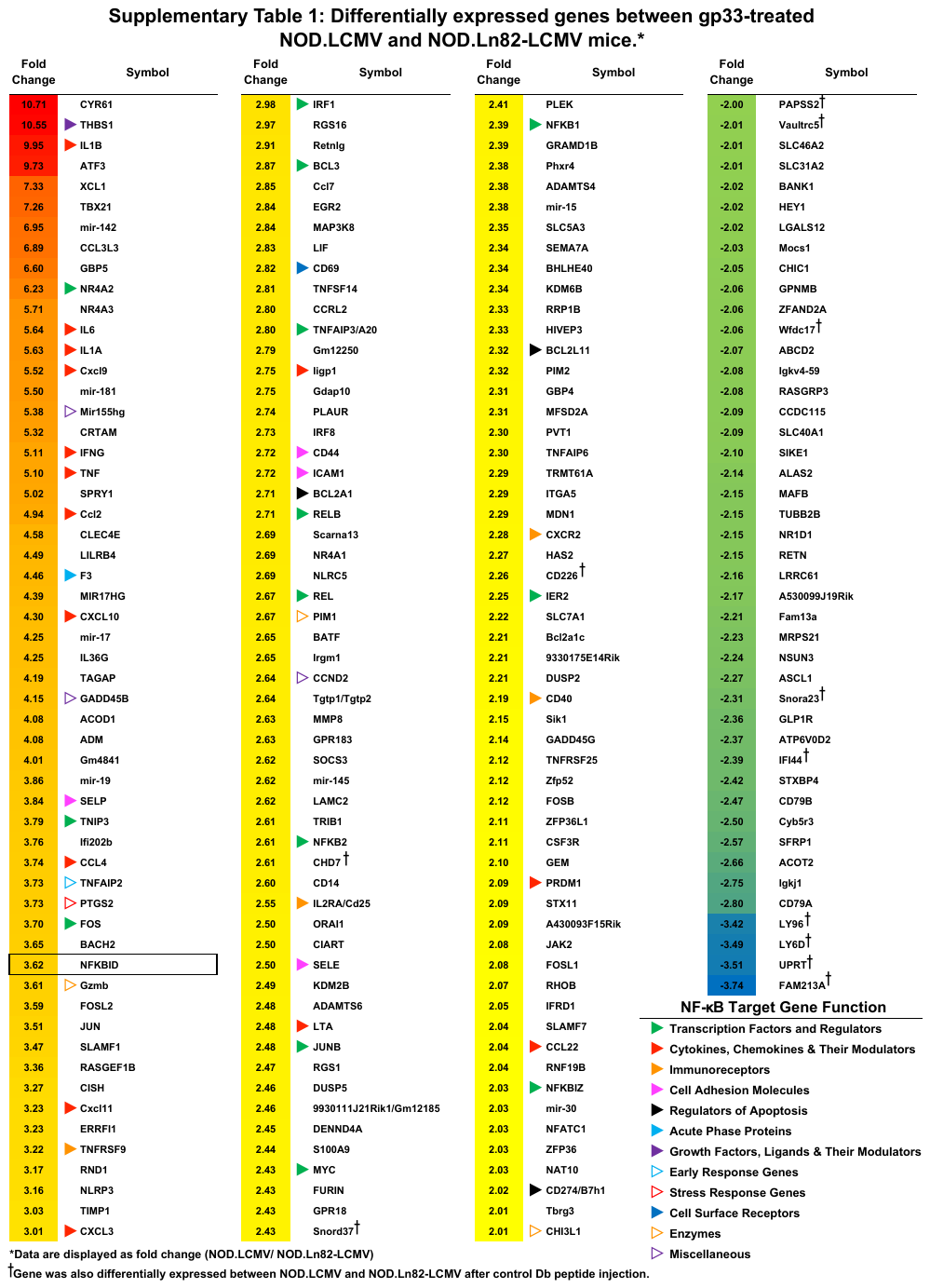

**Supplementary Table 2: qPCR primers for NF-κB target and NF-κB family member genes**

| **Gene** | **Forward** | **Reverse** |
| --- | --- | --- |
| *A20* | 5’- ACTCGGAAGCACCATGTTTG -3’ | 5’- CCTGTGTAGTTCGAGGCATGT -3’ |
| *Bcl2a1a* | 5’- ATACGGCAGAATGGAGGTTGG -3’ | 5’- TGGTCCGTAGTGTTACTTGAGG -3’ |
| *Bcl3* | 5’- CCGAATACTCAGCCTTTTCAAGC -3’ | 5’- TGGTAATGTGGTGATGACAGCC -3’ |
| *Cd69* | 5’- ACTGGAACATTGGATTGGGC -3’ | 5’- TTTTTGTGGTTCACGGACACG -3’ |
| *Gadd45b* | 5’- ATGAATGTGGACCCCGACAGC -3’ | 5’- CAGCAGAACGACTGGATCAGG -3’ |
| *Ier2* | 5’- GTGCTTCCTGTTGCCTTGC -3’ | 5’- TTGGTTCAGTTCAGGCTCACC -3’ |
| *Irf1* | 5’- GCAGATGGACATTATACCAGATAGC -3’ | 5’- CTCGTCTGTTGCGGCTTCG -3’ |
| *Nfkb1* | 5’- AGCACATAGATGAACTCCGGG -3’ | 5’- GATAGCAGTGGGCTGTCTCC -3’ |
| *Nfkb2* | 5’- GGGGACTTGGTGGGTCTATT -3’ | 5’- GCTCCCATAGGCACTGTCTT -3’ |
| *Nfkbia* | 5’- ATGAGGAGAGCTATGACACGG -3’ | 5’- TCCACATTCTTTTTGCCACTTTCC -3’ |
| *Nfkbid (Wt)* | 5’- TCCATGCTACCTATACAGAGGG -3’ | 5’- AGAGAGTCCTCCATCACCAGG -3’ |
| *Nfkbid (Total)* | 5’- CTGGGAGCAGAGCCTAACG -3’ | 5’- GGACTTAAACACAGCCGAAAGG -3’ |
| *Nfkbiz* | 5’- GAATCGGCAGTCTCATTCGC -3’ | 5’- GCCACTCTTACGATCCTTTGC -3’ |
| *Nr4a2* | 5’- GTTAAAGAAGTGGTTCGCACGG -3’ | 5’- AATCGACGTGGGCTCTGACG -3’ |
| *Rel* | 5’- CTGAGAAACCAAGAACTGCCCC -3’ | 5’- GAATAGTAAGGTTCAGCTTGCCC -3’ |
| *Rela* | 5’- ATCGAACAGCCGAAGCAACG -3’ | 5’- GCCATTGATCTTGATGGTGGG -3’ |
| *Relb* | 5’- GCCTTGGGTTCCAGTGACC -3’ | 5’- GCCTTGGGTTCCAGTGACC -3’ |
